# Mechanical control of the mammalian circadian clock via YAP/TAZ and TEAD

**DOI:** 10.1101/2022.02.04.478830

**Authors:** Juan F. Abenza, Leone Rossetti, Malèke Mouelhi, Javier Burgués, Ion Andreu, Keith Kennedy, Pere Roca-Cusachs, Santiago Marco, Jordi García-Ojalvo, Xavier Trepat

## Abstract

Circadian rhythms are a key survival mechanism that dictates biological activity according to the day-night cycle. In animals, cell-autonomous circadian clocks can be found in nearly every cell type and are subjected to multi-layered regulation. Although these peripheral clocks are remotely controlled by the master clock in the brain, they are also sensitive to their immediate mechano-chemical microenvironment. Whereas the mechanisms by which biochemical signalling controls the circadian clock at the single cell level are increasingly well understood, mechanisms underlying regulation by mechanical cues are still unknown. Here we show that the circadian clock in fibroblasts is regulated mechanically through YAP/TAZ and TEAD. We use high-throughput analysis of single-cell circadian rhythms and apply controlled mechanical, biochemical, and genetic perturbations to study the expression of the clock gene *Rev-erbα*. We observe that *Rev-erbα* circadian oscillations are disrupted concomitantly with the translocation of YAP/TAZ to the nucleus. By targeted mutations and tuning expression levels of YAP we identify TEAD as the transcriptional effector of this mechanosensitive regulatory pathway. Our findings establish a mechanism that links cell mechanobiology and the circadian clock, which could contribute to explain the circadian impairment observed in cancer and ageing, where the regulation of the mechanical environment and YAP/TAZ is lost.

## Introduction

The vast majority of organisms display daily physiological and behavioural changes in adaptation to the cyclic environment that arises from the Earth’s rotation ^1^. These rhythms, termed *circadian*, are endogenous and self-sustained at the cell level by a *clock* directed by set of proteins whose expression is regulated via a transcription-translation feedback loop with a period close to 24 hours ^12^. In mammals, up to 25% of the proteome is estimated to be influenced, in a cell type-specific manner, by the circadian clock, which deeply conditions dynamics and function of tissues and organs throughout the day ^13–15^.

The central element of the mammalian circadian clock is the transcription factor BMAL1:CLOCK, which activates the transcription of, among many others, *Cry* and *Per*. In turn, the sustained expression of CRY and PER generates a transcriptional loop by binding to BMAL1:CLOCK and blocking its activity ^16^. A second stabilizing layer of regulation of the clock is provided by REV-ERB, which, on one hand, is transcriptionally promoted by BMAL1:CLOCK and, on the other hand, acts as a transcriptional inhibitor of *Bmal1* ^7^.

For every cell residing in a peripheral tissue, the robustness of the circadian clock and its synchrony with the environment are subjected to multi-layered regulation ^2,3^. Decades of work have unveiled the biochemical side of this regulatory network which, mainly through neurocrine and paracrine signalling, drives important changes in the gene expression landscape of the different cell types ^4,17,18^. Besides biochemical regulation, recent evidence suggests a reciprocal communication between the circadian clock and mechanobiological factors such as actin dynamics, cell-cell adhesions and the stiffness of the extracellular matrix (ECM) ^5,6,19,20^. However, how and to what extent mechanotransduction impacts on the transcriptional regulation of the circadian network at a single cell level is still unknown.

By combining confocal microscopy, microfabrication, and a customized computational toolkit, we show that high nuclear levels of the transcriptional regulators YAP/TAZ cause a dramatic change in the expression of *Rev-erbα* and the subsequent malfunction of the fibroblast circadian clock. This novel molecular mechanism, which we show to be TEAD-dependent, could explain the highly impaired circadian clock in circumstances typically associated with high nuclear YAP/TAZ translocation such as cancer or ageing ^8–11^.

## Results

### *Rev-erbα* basal expression and circadian oscillations depend on cell density

We conducted our work using NIH3T3 fibroblasts, a cellular model extensively reported to express a robust self-sustained circadian molecular clock ^21,22^. We generated a stable clonal cell line carrying both a reporter of *Rev-erbα* transcription (RevVNP) ^21^ and a chimeric histone2B-mCherry that we used as a constitutive nuclear marker (Figure 1A, left). To avoid the effect of specific signalling pathways derived from hormonal shocks ^23^ and reveal underlying mechanobiological signals, cells were not synchronized by exogenous entrainment during our experiments. As a consequence of this, instead of performing the common analysis of populational oscillations, we systematically tracked and measured the single cell expression of RevVNP, which was obtained over the course of 3-day time-lapses performed by confocal microscopy. We then developed a method to quantify and compare to which extent the unsynchronised cells display a 24-hour oscillatory behaviour. Performing Fourier analysis of the fluorescence emission of each cell we obtained individual power spectral densities. From these we quantified the circadian power fraction, defined as the fraction of power found in a window between 0.7 and 1.3 days^-1^ of the cell’s RevVNP signal (Figure 1A, Figure S1, Materials and Methods). This analysis allows us to quantitatively distinguish between conditions that result in robust circadian oscillations (circadian power fraction tending to 1) from conditions where the RevVNP signal is affected by non-circadian modes, including non-rhythmic contributions (circadian power fraction tending to 0).

**Figure 1.**
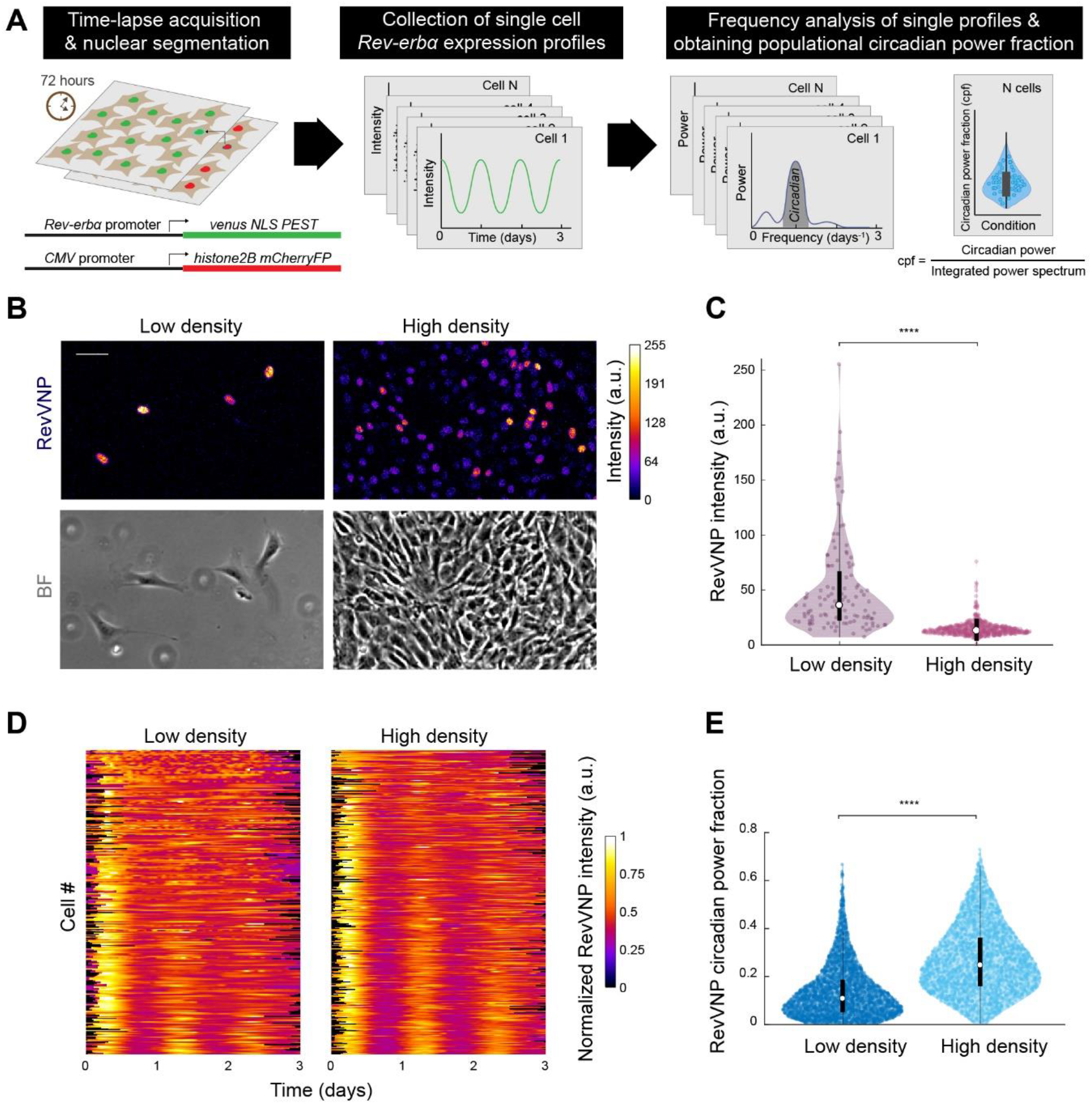
*Rev-erbα* basal expression and circadian oscillations depend on cell density. (A) Schematics of the systematic computational analysis pipeline used to calculate the circadian power fraction of a population of cells expressing RevVNP and H2B-mCherry and imaged during 72 hours via time-lapse confocal microscopy. (B) Confocal microscopy (top) and phase contrast (bottom) images of RevVNP-expressing cells grown at low (left) and high (right) density. Scale bar, 50 µm. (C) Violin plots representing the distribution of the single-cell RevVNP intensities of low- and high-density populations of a prototypic experiment of 14; n = 112 cells and n=622 for low density and high density, respectively; Medians and interquartile ranges are depicted as white circles and black bars, respectively. Two-sided Wilcoxon rank sum test; **** indicates a p-value < 0.001. Full p-values are reported in Table S1. (D) Raw data of the RevVNP intensities over time, represented in kymograph style, of 325 cells grown and tracked under low- or high-density conditions, from a representative experiment of 14. The single tracks are ordered from less (top) to more (bottom) circadian power fraction and aligned along the time axis according to maximum cross-correlation with the median track. (E)Violin plots representing the distribution of the single-cell RevVNP circadian power fraction of low- and high-density populations; n = 2726 and 2389 cells, respectively, from four experiments; Medians and interquartile ranges are depicted as white circles and black bars, respectively. Two-sided Wilcoxon rank sum test; **** indicates a p-value < 0.001. Full p-values are reported in Table S1.

We first addressed the effect of cell density on RevVNP expression by culturing cells on hydrogels (30 kPa in stiffness) at two distinct densities (∼30 cells/mm^2^ and 1000 cells/mm^2^, hereafter referred to as low and high density, respectively), which we kept approximately constant over time by arresting the cell cycle with the addition of thymidine, a DNA synthesis inhibitor ^24^. We observed a dramatic cell density-dependent difference in *Rev-erbα* expression, with cells cultured at low density displaying much higher RevVNP nuclear intensity than confluent cells (Figure 1B-C). These differences were ratified at the protein level via immunostainings against REV-ERBα (Figure S2). The single cell frequency analysis of the RevVNP signal revealed that the population of cells grown at high density mostly showed stable oscillations and had a higher circadian power fraction with an average period of 24.7 ± 1.9 h (Figure 1D-E). By contrast, cells at low density displayed low RevVNP circadian power fraction, with many cells exhibiting poor circadian oscillations, missing one or several peaks or even experiencing non-periodic fluctuations rather than oscillatory behaviour (Figure 1D-E).

### *Rev-erbα* circadian oscillations do not depend on cell-cell adhesions or paracrine signalling

We next asked whether the detected differences in circadian power fraction are caused by paracrine signals, as previously reported for intercellular circadian coupling and synchronization ^25,26^. To address this question, we exposed closely packed cells to an abrupt drop in cell density. To do so, two cell monolayers were grown to confluence separated by a PDMS barrier. Upon lifting the barrier (Figure 2A, left), cells at the monolayer edges begun to spread and migrate towards the freely available gap. These cells experienced a sudden increase in RevVNP intensity that transiently disrupted their circadian oscillations until the gap was closed (Figure 2A-C; Video S1). This disruption contrasted with the behaviour of cells far from the edge, which oscillated robustly throughout the entire experiment as shown in the kymograph depicted in Figure 2B. We obtained similar results in presence or absence of thymidine (Figure S3), which confirms that the increase seen in *Rev-erbα* transcription is not a side effect of a cell cycle re-entry at the monolayer edge upon a loss of cell-cell contact. These experiments further demonstrate that cell density regulates the circadian clock and suggest that this phenomenon is not explained by paracrine signals, as high- and low-density cells shared the same extracellular chemical environment. We confirmed that changes in paracrine signalling are not the cause of RevVNP circadian impairment by growing cells at low density with conditioned medium obtained from a high-density culture. We observed no increase in circadian power fraction compared to the low-density cells grown in fresh medium (Figure 2D). These results suggest that in our system cell density does not affect the circadian expression of *Rev-erbα* through paracrine signals, which led us to hypothesize that the observed behaviour had a mechanochemical origin.

**Figure 2.**
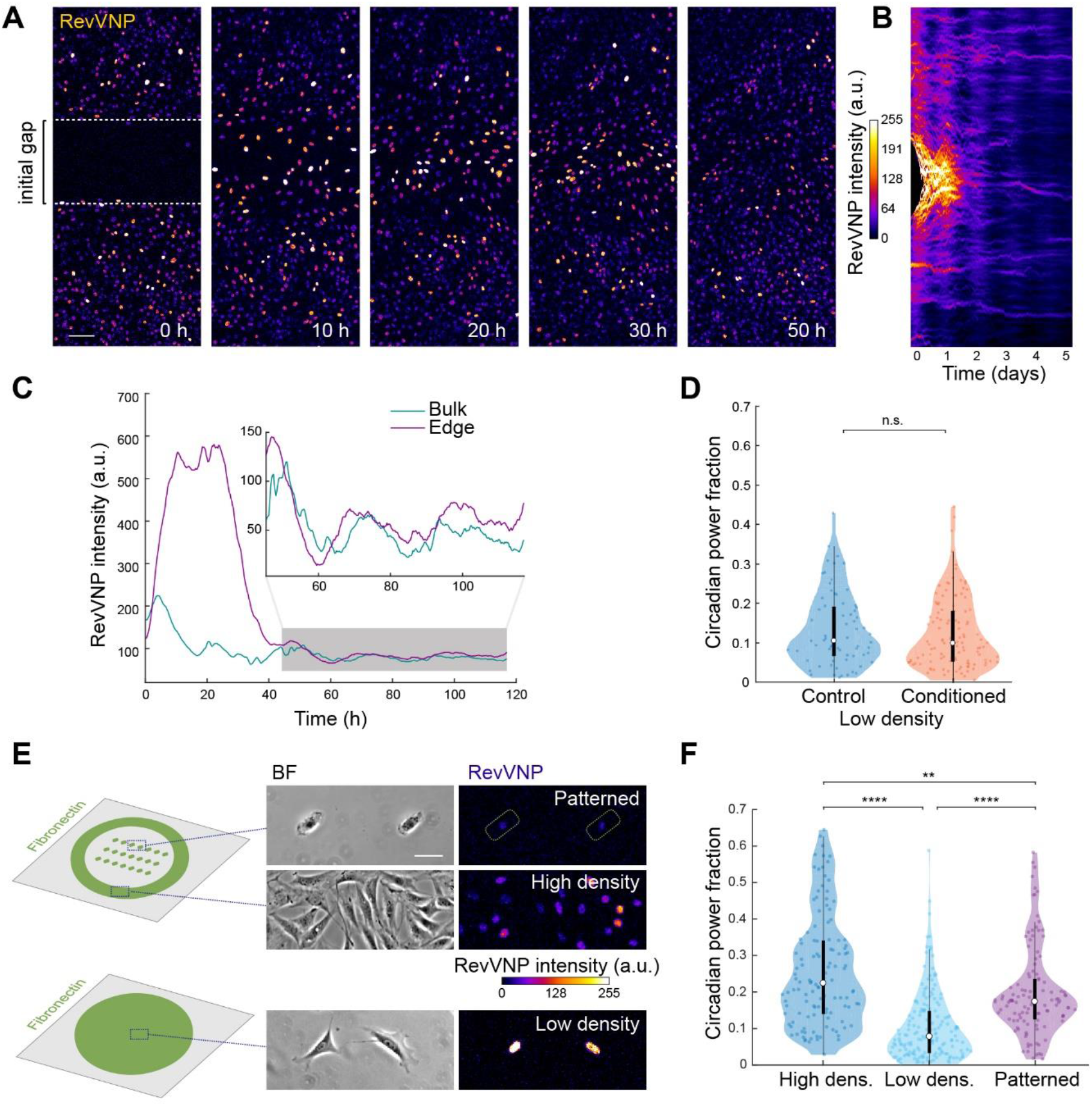
*Rev-erbα* circadian oscillations do not depend on cell-cell adhesions or paracrine signalling. (A) RevVNP-expressing cells were cultured in two adjacent compartments separated by a barrier until they reached high density. Next, the barrier was removed, and time-lapse confocal microscopy was performed during and after gap closure every 15 minutes for 5 days. A sequence of time-lapse images corresponding to the closure of the gap is shown. The prior location of the barrier is depicted in the left image as a white dashed line. (B) Kymograph representing the average RevVNP intensity per time point of all the cells from (A) along the x axis. (C) Average RevVNP intensity over time of the cells at the edge in comparison to those residing in the confluent zone. The inset shows a magnification of the area shaded in grey. This figure shows an example of n = 6 independent experiments. Scale bar, 100 µm. (D) Violin plots representing the distribution of the single-cell RevVNP circadian power fractions of low-density cells grown in fresh medium and conditioned medium; n = 72 and 116 cells, respectively, from three experiments; Medians and interquartile ranges are depicted as white circles and black bars, respectively; Two-sided Wilcoxon rank sum test; p-value = 0.4168. (E) Schematics of the strategy followed to isolate single cells via micropatterning of fibronectin on glass. On the right, phase contrast and confocal microscopy images of cells in stadium-shaped patterns (top), confluent cells cultured in the same well (middle), and non-confined cells cultured on a homogenous fibronectin-coated surface in a density as low as that of the micropatterned cells (bottom). Scale bar, 50 µm. (F) Violin plots representing the distribution of the single-cell RevVNP circadian power fractions of the conditions depicted in (E); n = 121, 174 and 115 cells for the high-density, low-density and micropatterned cells, respectively, from three experiments; Medians and interquartile ranges are depicted as white circles and black bars, respectively. Two-sided Wilcoxon rank sum test; ** indicates p-value < 0.01; **** indicates p-values < 0.0001. Full p-values are reported in Table S1.

To test this hypothesis, we first asked whether the differences in RevVNP circadian oscillations between high and low density depend on cell-cell adhesion. To address this question, we confined individual cells in fibronectin micropatterns of 1200 µm^2^ (a size comparable to that of fibroblasts within a monolayer) (Figure 2E). The confined individual cells showed a significant reduction in RevVNP intensity together with an increase in circadian power fraction when compared to free cells grown at equivalent low density, a behaviour resembling that of high-density cells (Figure 2E-F; Video S2). This result implies that contact-based mechanisms in general, and intercellular adhesion in particular, do not explain the dependence of the *Rev-erbα* circadian oscillations on cell density.

### *Rev-erbα* circadian power fraction anticorrelates with nuclear YAP

We thus explored whether the onset of circadian oscillations at high density had a contact-independent mechanical origin. We focussed on two key mechanosensitive transcriptional regulators. The first one is MAL, whose localization is regulated by actin dynamics and has been shown to modulate the transcription of the circadian gene *Per2* ^20^. The second one is YAP, the central component of the Hippo pathway, which translocates in a mechanoresponsive manner between the cytosol and the nucleus ^27–30^. YAP was recently shown to bind to REV-ERBα ^31^ although it has not been linked to the circadian clock. We measured and compared both circadian power fraction and subcellular localization of MAL and YAP in a series of conditions known to impact their nucleocytoplasmic shuttling. These conditions include the already mentioned high and low cell density and single cell micropatterning; treatment with drugs that alter the actomyosin cytoskeleton such as jasplakinolide, latrunculin A, para-nitro-blebbistatin and cytochalasin D; and the growth of cells on very soft hydrogels (300 Pa) ^32–35^. As expected, these conditions had a variety of effects on cell morphology (Figure 3A, Figure S4). We found no correlation between nuclear to cytoplasmic ratio of MAL and circadian power fraction (Figure S4). By contrast, circadian power fraction showed a progressive decrease with YAP nuclear to cytoplasmic ratio (Figure 3). This striking anticorrelation reveals that the robustness of the *Rev-erbα* circadian expression depends on the nucleocytoplasmic transport of YAP and its mechanosensitive regulation.

**Figure 3.**
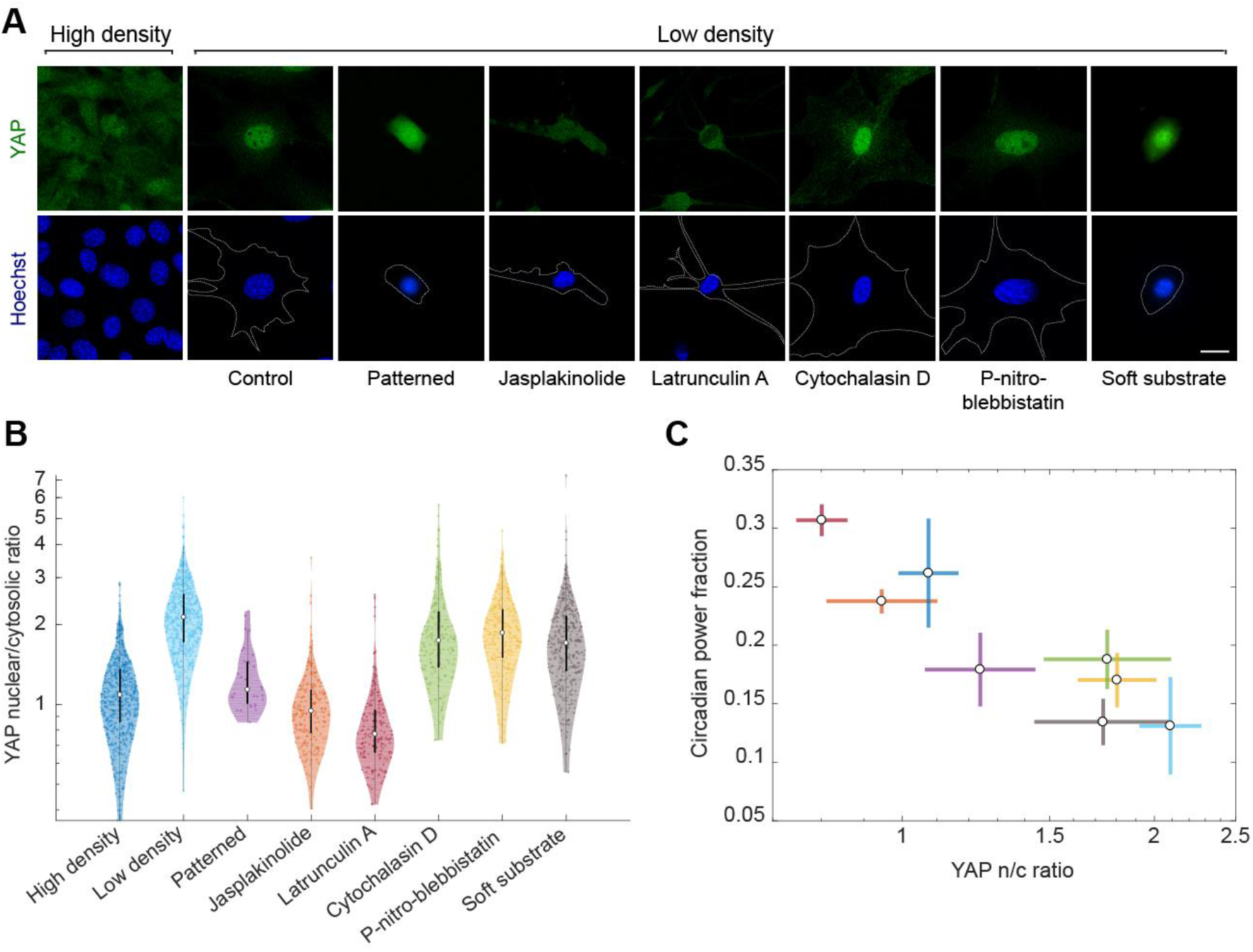
*Rev-erbα* circadian power fraction anticorrelates with nuclear YAP. (A) Confocal microscopy images of cells under different conditions (high-density control, low-density control, micropatterned cells, low-density treated for 24 hours with jasplakinolide 1 µM, latrunculin A 200 nM, cytochalasin D 1 µM, para-nitro-blebbistatin 10 µM and low-density cells grown on polyacrylamide gels with a stiffness of 300 Pa). The cells were stained with an anti-YAP antibody (green, top) and Hoechst (blue, bottom; the cell perimeter is represented with a dashed white line. Scale bar, 20 µm. (B) Violin plots representing the distribution of the single-cell YAP nuclear to cytosolic ratios for the conditions depicted in (A); n= 554, 589, 31, 203, 171, 238, 153 and 315 cells, respectively, from 3 to 7 experiments depending on the condition. Medians and interquartile ranges are depicted as white circles and black bars, respectively. (C) Correlation between circadian power fraction and YAP nuclear to cytosolic ratio of all the aforementioned conditions. The values represented are the means of the medians of each independent experiment for every condition. The error bars refer to the corresponding standard deviations. Pearson’s correlation coefficient is r = -0.89 with a p-value of 0.002.

### YAP/TAZ perturbs the circadian clock via TEAD

To test whether this dependence is causal, we induced in high-density cells a sustained overexpression of a non-phosphorylatable YAP mutant, 5SA-YAP, which has abnormally high nuclear retention ^36^. The resulting high levels of 5SA-YAP provoked a severe reduction of RevVNP circadian power fraction (Figure 4A-B) and an increase in REV-ERBα basal levels when compared to high-density control cells (Figure 4C-E). A similar result was obtained when overexpressing the TAZ mutant 4SA-TAZ, which is also retained in the nucleus due to the mutation of its four Lats inhibitory phosphorylation sites ^37,38^ (Figure S5). Together, these results establish that the circadian clock is controlled through both YAP and TAZ.

**Figure 4.**
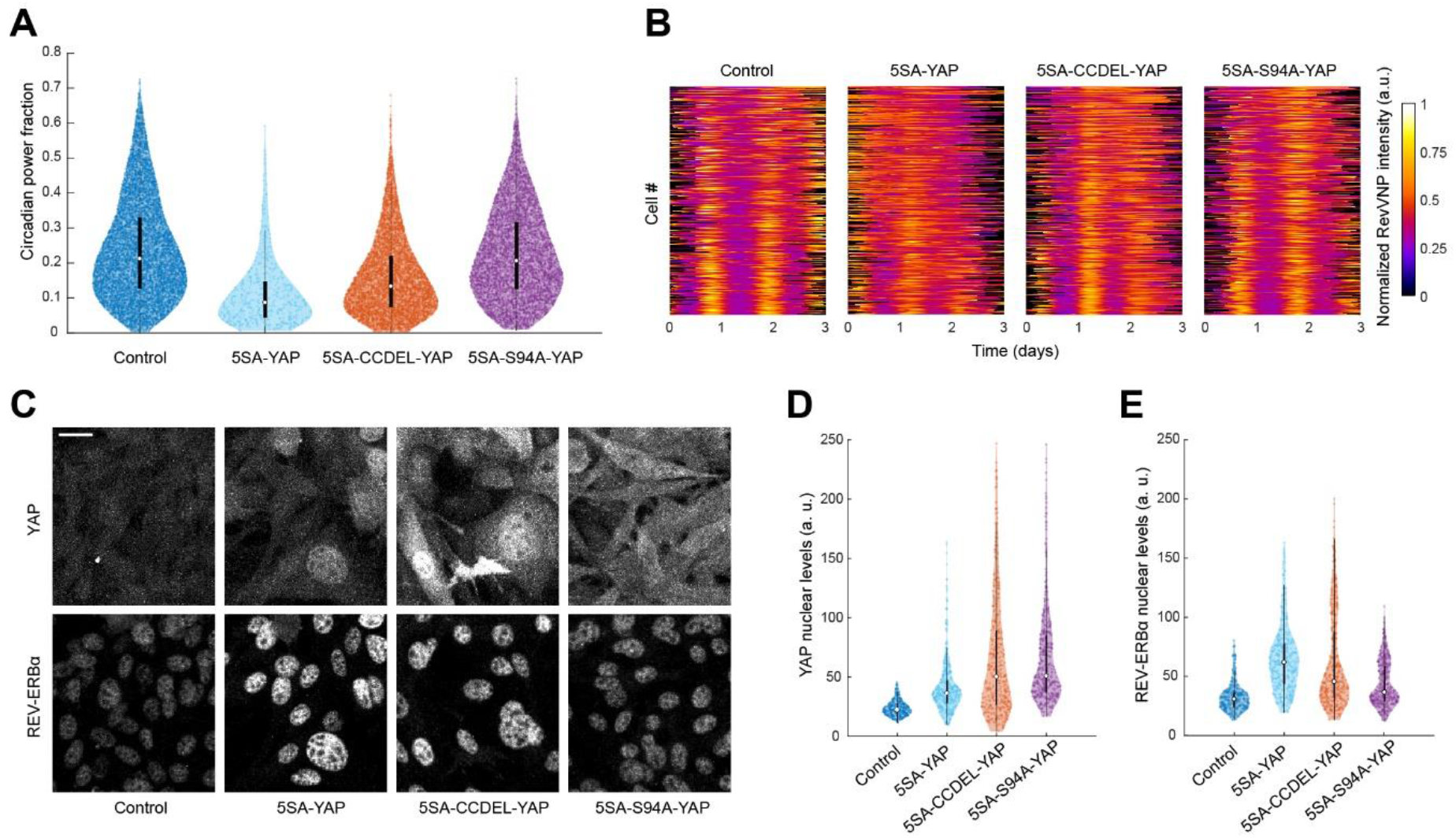
YAP perturbs the circadian clock via TEAD. (A) Violin plots representing the distribution of the single-cell RevVNP circadian power fractions of YAP-overexpressing cells carrying different mutations and the control; n = 6466, 440, 5320 and 4571 for control, 5SA-YAP, 5SA-CCDEL-YAP, and 5SA-S94A-YAP, respectively, from five independent experiments; Medians and interquartile ranges are depicted as white circles and black bars, respectively. Full p-values are reported in Table S1. (B) Confocal microscopy images of control, 5SA-YAP, 5SA-CCDEL-YAP, and 5SA-S94A-YAP cells immunostained for YAP (left) and a REV-ERBα (right). All images are displayed under the same brightness and contrast settings. Scale bar, 20 µm. (C-D) Violin plots representing the single-cell levels of nuclear YAP (C) and nuclear REV-ERBα (D) for all the conditions depicted in (B). n = 294, 400, 383 and 393 for control, 5SA-YAP, 5SA-CCDEL-YAP, and 5SA-S94A-YAP, respectively, from three independent experiments. Medians and interquartile ranges are depicted as white circles and black bars, respectively. Full p-values are reported in Table S1.

The activity of YAP/TAZ as gene expression coactivators requires the interaction with other transcription factors. Since YAP interacts with REV-ERBα through its coiled-coil (CC) domain ^31^, we tested if this interaction gives rise to the deleterious effect of nuclear YAP on the circadian clock. For that, we overexpressed 5SA-YAP but with its CC domain deleted (5SA-CCDEL-YAP), a mutation that had been previously shown to disable the molecular interaction between YAP and REV-ERBα ^31^. We found that the CCDEL mutation only rescued slightly the impaired RevVNP circadian oscillations (Figure 4A) and the REV-ERBα basal levels (Figure 4B-D) observed upon 5SA-YAP overexpression. This result shows that the direct molecular binding through the YAP CC domain is not the main regulatory pathway connecting YAP and the circadian clock.

Alternatively, we studied whether this regulation was mediated by the well-known interaction of YAP/TAZ with the TEAD family ^39^. Analysis of published ChIP-seq and microarray databases revealed that *Bmal1* -but not *Rev-erbα*- is listed as a YAP and TEAD target ^40–43^, providing a potential direct mechanism for YAP/TAZ to control the circadian clock. To test this possibility, we overexpressed 5SA-S94A-YAP, a mutant version of YAP unable to interact with TEAD ^42^ and observed that the cells recovered, to a large extent, both the RevVNP circadian power fraction and the REV-ERBα basal levels displayed by the wild-type high-density population (Figure 4). This result establishes that the circadian clock deregulation in nuclear YAP-enriched cells is caused by the YAP-TEAD transcriptional cascade.

## Discussion

The blooming of mechanobiology in this century has established mechanics as an essential player in the regulation of key biological processes in physiology and disease ^44^. However, it was not until very recently that the influence of mechanical cues on the circadian clock was addressed, partly due to large differences in experimental approaches and timescales between the fields of chronobiology and cell mechanics. Traditionally, circadian experiments have mainly been carried out at the populational level and, when performed in vitro, they often relied on cell synchronisation by the application of *zeitgebers* ^20,23,45^. These treatments are biochemically invasive and limit the in-depth study of non-canonical signalling cascades like those triggered by mechanical factors, as well as the single cell circadian regulation in heterogeneous and complex environments. Here we studied the mechanical regulation of the circadian clock at the single cell level and without need for synchronization strategies. This approach allowed us, for example, to study oscillations in single patterned cells and to closely link cell density with the oscillation robustness. We expect the present work to open new avenues of research to address how mechanosensing signalling cascades impact the clock. This approach can be fostered by a) the development of techniques allowing a precise mechanical manipulation of the cells; b) the increasing availability of new fluorescent circadian reporters through genome-editing techniques like CRISPR knock-in ^46,47^ and c) the continuous development of new algorithms to analyse noisy oscillatory signals in single cells ^48^.

An intricate relationship between cellular mechanobiology and the circadian clock has recently begun to emerge. For example, two recent studies have shown that actin dynamics and the secretion of extracellular matrix are subjected to circadian regulation ^6,49^. The groups of C. H. Streuli and Q. J. Meng have studied the complementary regulation, showing that the robustness of the circadian clock is influenced by the stiffness of the extracellular environment, with a mechanistic focus on integrins and Rho/ROCK ^19^. With our finding that nuclear YAP/TAZ disrupts Rev-erbα oscillations via TEAD, we now provide a transcriptional explanation of how mechanical signals, independently of their nature and how they are triggered, can be transduced into alterations of the circadian clock. Given the widespread role of YAP/TAZ as core mechanosensors, the potential implications of this new mechanism are multiple. For example, we believe this study will help contribute to a more complete understanding of the loss of circadian rhythms in ageing and cancer, two processes associated with an impaired clock, an alteration in the mechanical properties of the tissues and an abnormal activation of YAP/TAZ ^8,11,28,50–54^.

Finally, our result that YAP/TAZ and TEAD dramatically affect the basal expression of REV-ERBα have potential implications beyond the field of chronobiology. REV-ERB proteins are nuclear receptors that, acting as strong transcriptional repressors, play a key role in a large variety of physiological processes, and have been extensively used as drug targets ^55,56^. For example, high levels of REV-ERB have been correlated to increased insulin secretion and to altered lipid metabolism, which has opened new strategies for the treatment of diabetes ^56^. REV-ERB proteins are also involved in inflammation regulation, muscle function through the control of mitochondrial activity, bone loss and even Alzheimer’s disease ^56^. In light of this, we expect this study to serve as a connection point between the largely disconnected fields of chronobiology, mechanobiology and metabolism.

## Materials and Methods

### Cell culture

We cultured NIH3T3 mice fibroblasts in Dulbecco’s Modified Eagle Medium (DMEM) with pyruvate (41966-029, ThermoFisher) supplemented with 10% or 2% FBS (for cell lines maintenance or for experimental purposes, respectively; 10270-106, ThermoFisher), 100 U ml^−1^ penicillin, 100 μg ml^−1^ streptomycin and 292 μg ml^−1^ of L-glutamine (10378016, ThermoFisher). For all the experiments except the gap closure one, thymidine 2 mM (T9250-1G, Sigma) was added 24 hours before fixation or image acquisition. The experiment with conditioned medium was carried out with medium composed by 50% collected medium from a high-density population of cells after two days of culture and 50% fresh medium. To prevent undesired entrainments caused by medium changes like described in ^57^, we maintained the same medium for the cell culture on the plates and the entire image acquisition. However, it was unavoidable the conveyance of a certain level of phase synchrony in RevVNP expression in some cases, mainly in the gap closure experiments, which required intensive cell culture manipulation.

### Cell lines and constructs

For the circadian analyses we generated a stable cell line expressing venus fluorescent protein with a nuclear localization signal (NLS) and a destabilization PEST domain under the promoter of Rev-erbα. We used specifically the plasmid rev12-ex2-Venus-NLS-PEST1, courtesy of Prof. Ueli Schibler (Nagoshi et al., 2004), and random genomic integration via, first, lipofectamine-based transfection (Lipofectamine 3000, ThermoFisher) and, second, several rounds of fluorescence-based flow cytometry sorting, until stable cassette expression was observed. Lentiviral particles carrying plenti6-H2B-mCherry (Addgene plasmid #89766 ^58^) were generated by transient transfection of Hek293T cells together with the corresponding envelope and packaging vectors. Several clonal lines were obtained by transduction of the RevVNP cell line with the mentioned particles and flow cytometry sorting (Beckman Coulter). For reproducibility reasons, one of them was selected and used for all the experiments.

The 5SA-YAP, 5SA-CCDEL-YAP, 5SA-S94A-YAP, 4SA-TAZ, and control cell lines were obtained by transduction with retroviral particles generated in Hek293T cells and sustained selection with puromycin 2 μg ml^-1^ (A1113803, ThermoFisher). Specifically, the retroviral FLAG-5SA-YAP and FLAG-5SA-S94A-YAP plasmids were generated via restriction-based cloning from pQCXIH-Myc-YAP-5SA (Addgene plasmid #33093 ^36^ and pCMV-Flag-YAP-5SA/S94A (Addgene plasmid #33103 ^42^, respectively, and pBABE-PURO (Addgene plasmid #1764 ^59^). The CCDEL deletion was introduced in the latter via overlapping PCR with the primers YAP-DEL-CC-HA-F and YAP-DEL-CC-HA-R as described in ^31^. The FLAG-5SA-S94A-YAP plasmid corresponds to the Addgene plasmid The FLAG-4SA-TAZ plasmid was a kind gift of Dr. Hyun Woo Park (Yonsei University, Seoul, Korea ^37^). All the cell lines were checked by PCR, DNA sequencing and Western blot.

The cytoskeletal drugs used throughout the study were: jasplakinolide 1 µM (J4580-100UG, Sigma), cytochalasin D 1 µM (C8273-1MG, Sigma), latrunculin A 200 nM (L5163-100UG, Sigma) and para-nitro-blebbistatin 10 µM (DR-N-111, OptoPharma). The control conditions were performed with similar concentrations of DMSO (D8418-50ML, Sigma). All of them were added 24 hours before the initiation of the live imaging or fixation.

### Time-lapse imaging of circadian oscillations of NIH3T3 fibroblasts

Time-lapse acquisitions were performed on a scanning confocal microscope (Nikon Ti Eclipse) equipped with thermal, CO2, and humidity regulation. 12-well or 6-well glass-bottomed plates were fixed on the stage and imaging was performed using a 10× objective (NA = 0.30, air) with focusing maintained by the Perfect Focus System (Nikon). The software NIS Elements was used to image every 15 minutes during experiments lasting for 3 to 5 days. Multiple wells, and multiple positions within the different wells, were imaged by phase contrast and in laser scanning mode using a 488 nm and a 561 nm excitation. For the 488 nm acquisition two different sets of settings were used, to better capture the large variations in fluorescence signal seen in the RevVNPchannel: a low-intensity channel with low excitation power, and a high-intensity channel with 5 times higher excitation power. All confocal images were acquired by scanning 1024×1024 pixels on the objective’s field of view, yielding a pixel-size of 1.28 um. Pixel dwell times were adjusted independently in each experiment.

### Single-cell tracking

Tiff stacks of time-lapse microscopy images were analysed using Fiji ^60^. Fluorescent nuclei visible in the high-intensity channel were tracked in space and time using the FIJI plugin TrackMate ^61^. Within TrackMate, spots were identified using the LoG detector, after which the LAP tracker was used, with a maximum linking distance of 15 pixels and gap closing set to 2 frames. The resulting tracks were then filtered by length, keeping only those that lasted at least 60 hours. In the rare cases of cell-division, only one daughter cell was tracked. The tracking data was then used to additionally obtain tracks for the low-intensity channel. Finally, the position and intensity data for both channels were exported as .csv files.

### Frequency Analysis of datasets of single-cell circadian oscillations

All analyses were performed with custom software created using MATLAB R2017a (Figure 1—figure supplement 1). To calculate a cell’s circadian power fraction firstly any missing time-points in each single-cell time-intensity track were interpolated. After this, each track was denoised by a low-pass filter, its mean value was subtracted, and the track was divided by its standard deviation. Then the tracks were analysed individually to remove any dying cells: these were defined as intensity traces whose standard deviation in the final 24 hours is less than 5% of the standard deviation during the rest of the experiment. A fast Fourier transform (FFT) was then applied, and the power spectral density (PSD) of each track was generated using the periodogram. All the PSDs were averaged, and the ensemble circadian frequency *f*_*c*_ was identified as the coordinate of the peak in the interval [0.7, 1.3] days^-1^ in the average PSD. The single-cell PSDs were then individually analysed as follows: the integral of the PSD in the frequency interval W = [*f*_*c*_ – 0.2, *f*_*c*_ + 0.2] days^-1^ was divided by the full integral of that individual track (i.e., the total power), giving what we termed the circadian power fraction of a single-cell oscillation. Note that to set the width of W we used pooled experiments with cells at high-density, identified the ensemble circadian frequency *f*_*c*_ and measured the full width at half maximum of the corresponding peak.

### Kymography

The denoised and rescaled single-cell tracks were further analysed by pairwise cross-correlation, and the phases of the signals were obtained as the lag values giving maximum cross-correlation were obtained. By virtue of the periodicity of the signals, these lags were restricted to the interval [-1,1] days. Tracks were ordered by decreasing circadian power fraction and the phases (taken with respect to the median track) were used to shift (in time) the other tracks and thus artificially synchronize the dataset. These data were plotted as a kymograph of fluorescence intensity by time and circadian power fraction rank, with the highest circadian power fraction tracks at the top.

For the gap closure experiments, the nuclei in time-lapse fluorescence images were segmented using a combination of TrackMate’s spot detection algorithm and intensity thresholding. The kymographs were generated by averaging the intensity of the segmented nuclei in each frame along the direction parallel to the wound edge. These lines were then assembled into a kymograph representing fluorescence intensity as a function of time and distance from the centre of the wound.

### Polyacrylamide gels

Polyacrylamide (PAA) gels with Young modulus E= 30 kPa – which matches approximately the stiffness of the majority of abdominal organs and the skin ^62^ - were produced according to a previously published protocol ^63^. A PBS solution containing 12% acrylamide (161-0140, Bio-Rad), 0.15% bis-acrylamide (161-0142, Bio-Rad); plus 0.05% ammonium persulphate (A3678, Sigma) and 0.05% tetramethyl ethylenediamine (T9281-25ML, Sigma) was prepared and allowed to polymerize between a coverslip and a glass-bottomed dish (Mattek). Alternatively, 3% acrylamide and 0.03% bis-acrylamide were used to make 300 Pa gels. The PAA gel surface was then incubated with a solution of 2 mg ml^−1^ Sulpho-SANPAH (4822589, ThermoFisher) under ultraviolet light for 5 min (wavelength of 365 nm at 5 cm distance). After that, the excess Sulpho-SANPAH was removed by three consecutive 3 min washes with PBS. A solution of fibronectin 0.1 mg ml^−1^ (F0895, Sigma) was added on top of the gels and left overnight at 4°C.

### Gap closure

The gap closure experiments were performed following the protocol described in ^64^. Magnetic PDMS stencils consisting in two hollow regions separated by a barrier of 600-900 µm were obtained using a customized 3D-printed mould. The cured, autoclaved, passivated, and dried magnetic PDMS stencils were then deposited on a fibronectin-coated PAA gel polymerized in a glass-bottomed dish. After this, the dish was placed on top of a custom-made magnetic holder to keep the gasket in place and avoid medium leakage. The hollow regions were then filled with 40,000 cells contained in 120 µl of medium. After 1 hour, excess of cells was removed by 2 consecutive washes with medium. The attached cells were left overnight at 37°C before the removal of the magnetic PDMS gasket and the filling of the whole glass-bottomed dish with medium.

### Immunostainings

Cells were fixed using paraformaldehyde 4% for 15 min and washed with PBS three times. Then, they were permeabilized with 0.1% Triton X-100 for 45 min, incubated with the primary antibodies for 90 min at room temperature, washed and incubated with the secondary antibodies for another 90 min at room temperature. After this step, Hoechst 33342 (H3570, ThermoFisher) was added at a concentration of 1 μg ml^−1^ for 10 minutes. Finally, cells were mounted in Mowiol reagent (81381, Merck). The buffers used during the whole procedure had fish gelatine 1.6% (v/v) as a blocking agent (G7765, Merck). The primary antibodies used were anti-NR1D1 (ab174309, abcam), anti-MKL1 (ab49311, abcam) and anti-YAP 63.7 (sc-101199, Santa Cruz). The secondary antibodies used were Alexa Fluor-488 anti-mouse (A-11029, ThermoFisher), Alexa Fluor-488 anti-rabbit (A-21206, ThermoFisher), Alexa Fluor-555 anti-mouse (A-21424, ThermoFisher) and Alexa Fluor-555 anti-rabbit (A-21429, ThermoFisher). All the antibodies were incubated in a 1:200 dilution. The acquisition of z-stacks (with a z-axis step of 0.7 µl) was done using a 40x water immersion LD LCI Plan Apo objective (NA = 1.2) in a scanning confocal microscope with Fast AiryScan (Zeiss LSM880) or a 60x oil immersion objective (NA = 1.40) in an inverted microscope (Nikon Eclipse Ti). Maximal projections of the stacks were obtained with Fiji before quantification of nuclear and cytosolic intensity levels or, for the case of YAP and MAL nuclear to cytosolic ratios, they calculated after measuring the mean intensity of two adjacent regions of identical size, inside and outside the nucleus as in ^34^ using the Hoechst image as a reference for the nuclear position.

### Micropatterning

The micropatterning protocol was an adaptation of ^65^. Single glass-bottomed dishes (35 mm Dish, No. 0 Coverslip, 10 mm Glass Diameter, MatTek) were cleaned with ethanol 96%, dried, and oxidized by exposure to oxygen-plasma (PCD-002-CE, Harrick) for 30 s at 7.2 W. After that, a ring of PDMS with an inner diameter of 7 mm was attached to the coverslip to delimitate the micropatterning area, and that area was immediately filled with 30 µl of PLL-PEG solution (PLL (20 Kda)-g[3,5]-PEG(2), SuSoS) at a concentration of 0.1 mg ml^−1^ in PBS, which was incubated for 1 h at room temperature. Then, the solution was carefully removed, the surface was washed three times with PBS and 30 µl of photoinitiator PLPP (4-benzoylbenzyl-trimethylammonium chloride; Alvéole) at a concentration of 14.5 mg ml^−1^ were added. After that, micropatterning was performed in an inverted microscope (Nikon Ti Eclipse), using the PRIMO system controlled by the Leonardo software (Alvéole) and a 20X objective (CFI S Plan Fluor ELWD ADM, Nikon), directing a UV power of 1050 mJ mm^-2^ following the patterns predesigned in ImageJ. When the patterning was completed, filtered PBS (concentration 1X) was used to clean three times and the PDMS ring was removed. The whole glass surface was then treated for 10 min with a solution of fibronectin (F0895, Sigma-Aldrich) 100 µg ml^-1^ and Alexa Fluor 647 fibrinogen (F35200, ThermoFisher) 5 µg ml^-1^, which caused attachment of the coating proteins at both the patterns and the area where the PDMS ring had previously been placed. After three washes, the glass-bottomed dishes were stored, filled with PBS, at 4°C for a maximum of 48 hours before the experiments. 2*10^5^ cells were seeded diluted in DMEM with 0.2% FBS to avoid cell attachment in unwanted areas. After one hour, excess of cells was removed by three intensive washes with the same medium. 24 hours before imaging or fixation the medium was replaced with 2% FBS-containing DMEM.

### Statistical analysis

All violin plots were generated using *violinplot*, an open-source MATLAB script ^66^. Two-sided Wilcoxon rank sum test were used when testing for statistical significance, and *p*-values smaller than 0.05 were considered as significant, unless specified otherwise.

## Data and code availability

The data that support the findings of this study and the MATLAB analysis procedures are available from the corresponding authors on reasonable request.

## Acknowledgments

We thank S. Aznar-Benitah, G. Solanas, M. Uroz, A. Elósegui-Artola, C. Pérez-González, D. Zalvidea and all the members of the Roca-Cusachs and Trepat laboratories for their discussions and support. We thank A. Menéndez, S. Usieto and N. Castro for daily technical assistance. We also thank N. Montserrat, E. Martínez, U. Schibler, H.W. Park and Y.S. Choi for sharing some of the cell lines and plasmids used in this work. This paper was funded by The Generalitat de Catalunya (AGAUR SGR-2017-01602 to X.T, AGAUR Beatriu de Pinós 2014 BP-B 00105 to J.F.A., the CERCA Programme, and “ICREA Academia” awards to P.R-C. and J.G.-O.); Spanish Ministry for Science and Innovation MICCINN/FEDER (PGC2018-099645-B-I00 to X.T., BFU2016-79916-P and PID2019-110298GB-I00 to P. R.-C, and PGC2018-101251-B-I00 to J.G.-O.); European Research Council (Adv-883739 to X.T.); Fundació la Marató de TV3 (project 201903-30-31-32 to X.T.); European Commission (H2020-FETPROACT-01-2016-731957 to P.R-C. and X.T., H2020 Marie Sklodowska-Curie Actions MECHADIAN - IF/750557 to J.F.A.); La Caixa Foundation (LCF/PR/HR20/52400004 to to P.R-C. and X.T.); IBEC is recipient of a Severo Ochoa Award of Excellence from the MINECO. DCEXS-UPF is recipient of a Maria de Maeztu Award of Excellence from the MINECO.

## Author contributions

J.F.A. and X.T. conceived the project. J.F.A., M.M., L.R. and I.A. performed experiments and analysed data. L.R. and J.B. developed software and analysed data. K.K., P.R.-C., S.M., and J.G.-O. contributed technical expertise, materials, and discussion. J.F.A., L.R. and X.T. wrote the manuscript. J.F.A. and X.T. supervised the project. All authors revised the completed manuscript.

