## Supplementary Material for "Mechanical control of the mammalian circadian clock via YAP/TAZ and TEAD"

### Supplemental information

**Video S1. The origin of the *Rev-erbα* circadian impairment at low density is mechanical rather than paracrine.** Time-lapse confocal imaging of RevVNP-expressing cells during and after gap closure. Cells were cultured on hydrogels at high density in two adjacent compartments separated by a PDMS barrier, which was removed just before imaging for 5 days every 15 minutes. Cells were not entrained by serum or hormonal shocks, although some synchronisation is observed, possibly arising from the unavoidable medium washes after seeding and after barrier removal. Scale bar, 100 µm.

**Video S2. Cell confinement in microprinted fibronectin areas rescues the *Rev-erbα* circadian impairment observed at low density.** Phase contrast and confocal microscopy time-lapse movies of cells confined in stadium-shaped fibronectin-coated patterns. Images were taken every 15 minutes for 3 days. Scale bar, 100 µm.

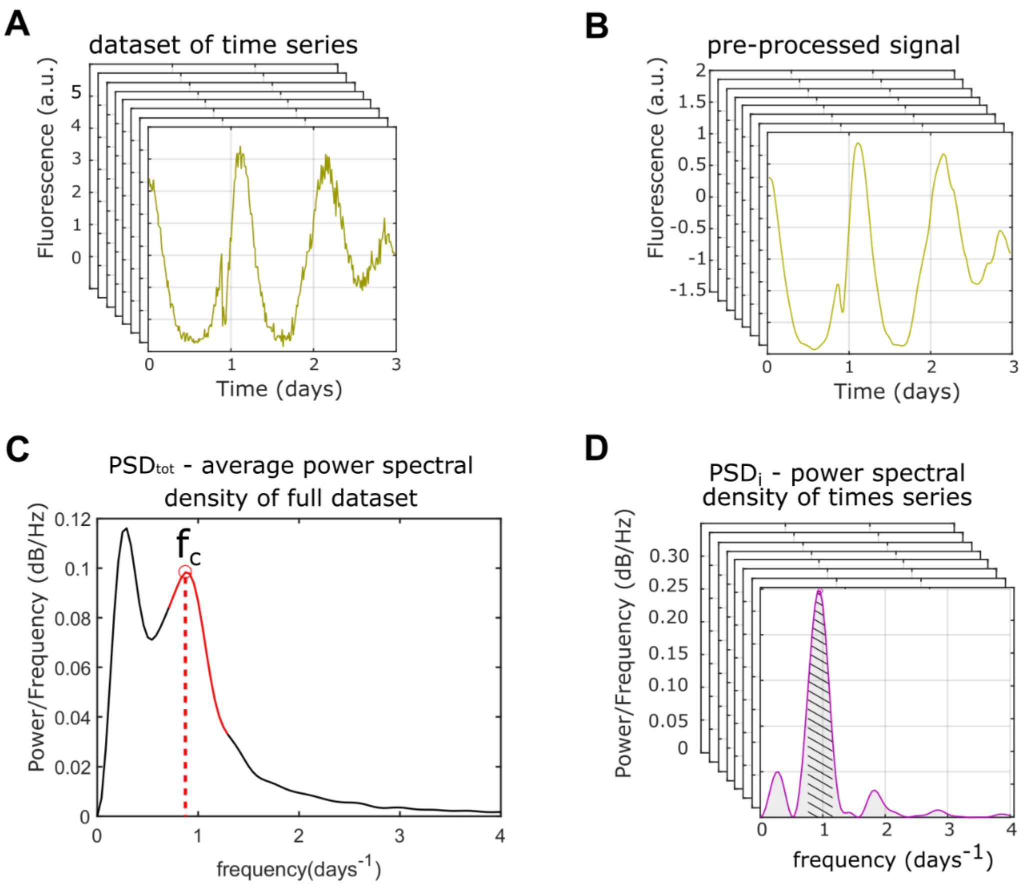


**Figure S1. Calculation of Circadian Power fraction from single cell RevVNP signals.**

(A) A dataset of single-cell RevVNP fluorescence signals is obtained from cell-tracking.

(B) the tracks are pre-processed by a subtracting the mean, dividing by the standard deviation and applying a low-pass filter.

(C) using MATLAB's fft function the individual power spectra $PSD_i$ of each track is calculated and averaged to give $PSD_{tot}$ from which the dataset's circadian frequency $f_c$ is extracted.

(D) each $PSD_i$ is integrated over an interval centred on $f_c$ (dashed area), and the result is normalized to the full integral (grey area) yielding a value of circadian power fraction between 0 and 1 for each track.

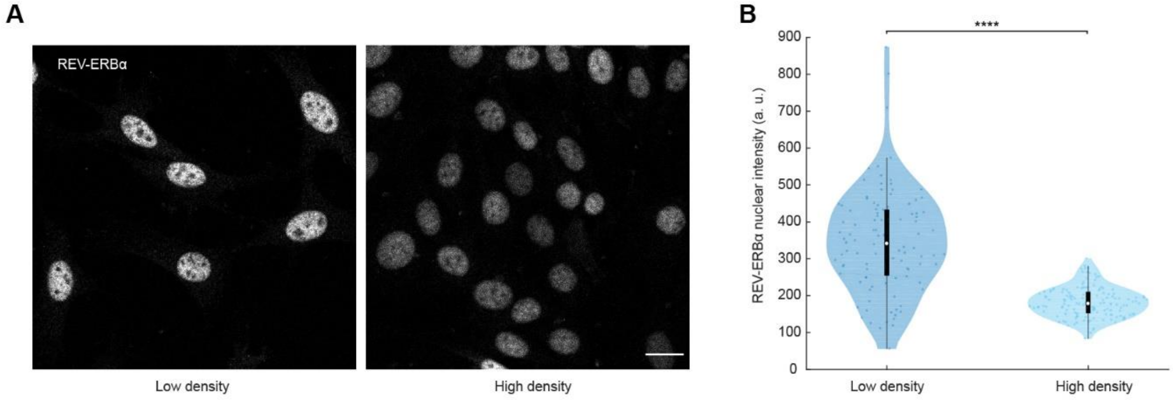


**Figure S2. REV-ERBα protein levels decrease with cell density.**

(A) Confocal microscopy images of low- and high-density cell populations immunostained against REV-ERBα. All images are displayed under the same brightness and contrast settings. Scale bar, 20 µm.

(B) Violin plots representing the single-cell levels of nuclear REV-ERBα. n = 104 for low density and 142 for high density, from one representative of five independent experiments. Two-sided Wilcoxon rank sum test; **** indicates p-value < 0.0001.

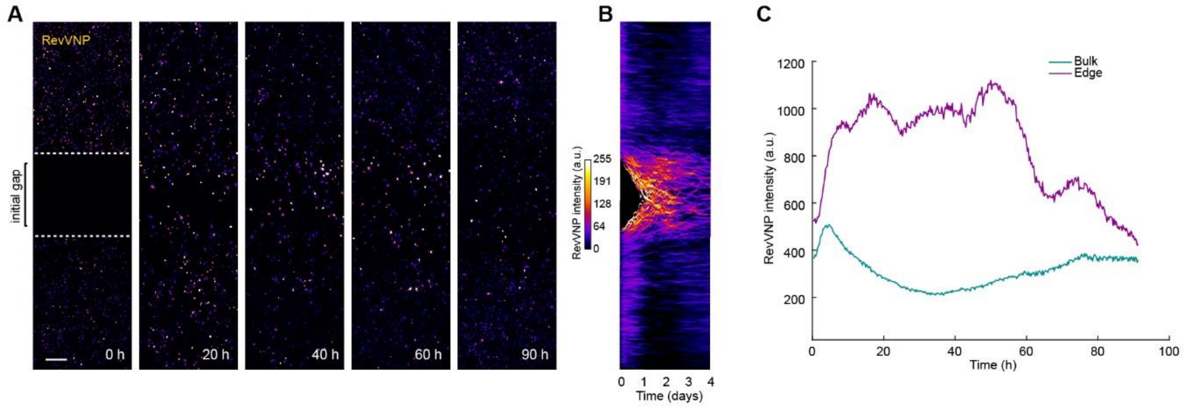


**Figure S3. The increase in *Rev-Erbα* expression upon transition from high to low density is not the consequence of cell-cycle re-entry**.

(A-C) RevVNP-expressing cells were cultured in two adjacent compartments separated by a barrier until they reached high density. Next, thymidine 2 mM was added, the barrier was released and time-lapse confocal microscopy was performed during and after gap closure every 15 minutes for 4 days. A selection of time-lapse images showing the closure of the gap is depicted in (A) while (B) is a kymograph representing the average RevVNP intensity of all the cells along the x axis per time point. The average RevVNP intensity over time of the cells at the edge in comparison to those residing in the confluent zone is shown in (C). The prior location of the barrier is depicted in the left image as a white dashed line. Scale bar, 100 µm.

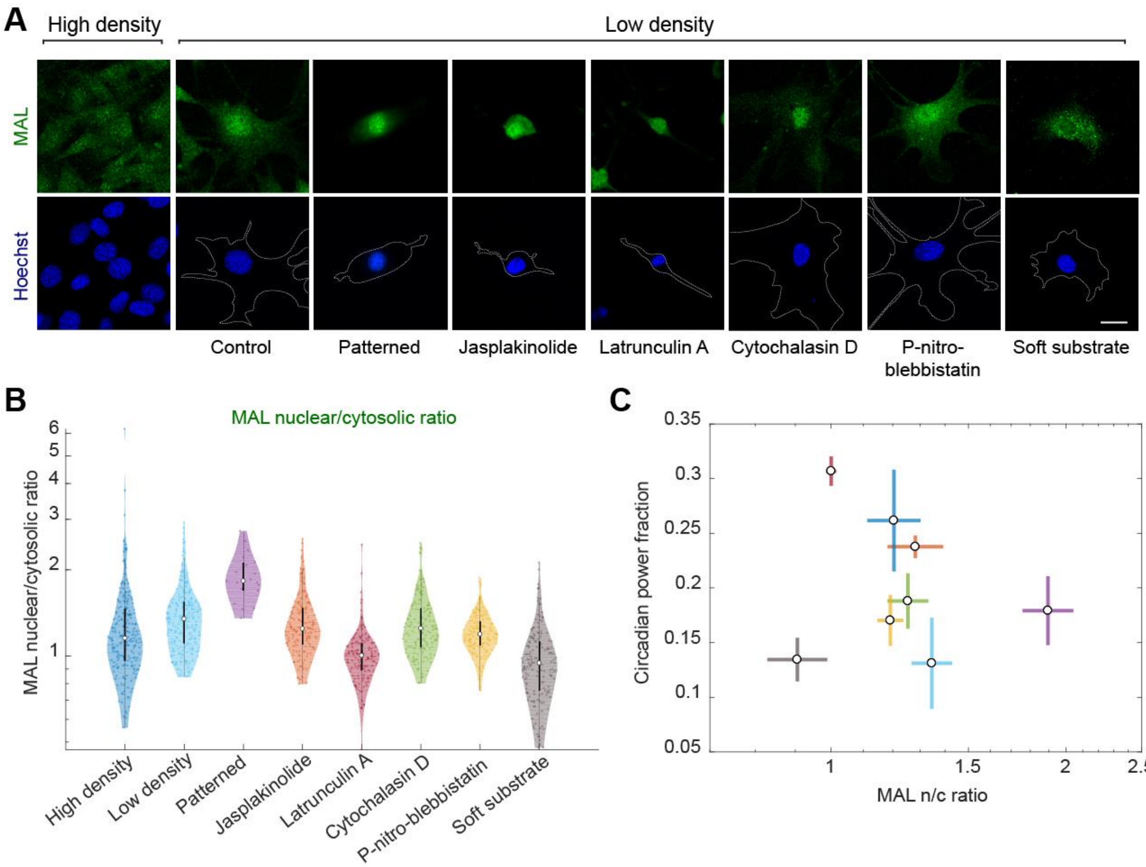


**Figure S4. *Rev-erbα* circadian power fraction does not correlate with nuclear MAL.**

(A) Confocal microscopy images of cells under different conditions (high-density control, low-density control, micropatterned cells, low-density treated for 24 hours with jasplakinolide 1 µM, latrunculin A 200 nM, cytochalasin D 1 µM, para-nitro-blebbistatin 10 µM and low-density cells grown on polyacrylamide gels with a stiffness of 300 Pa). The cells were stained with an anti-MAL antibody (green, top) and Hoechst (blue, bottom; the cell perimeter is represented with a dashed white line. Scale bar, 20 µm.

(B) Violin plots representing the distribution of the single-cell MAL nuclear to cytosolic ratios of the conditions depicted in (A); n = 215, 254, 19, 163, 140, 165, 157 and 162, respectively, from 3 to 4 experiments depending on the condition. Medians and interquartile ranges are depicted as white circles and black bars, respectively.

(C) Dependence of circadian power fraction with MAL nuclear to cytosolic ratio of all the aforementioned conditions. The values represented are the means of the medians of each independent experiment for every condition. The error bars refer to the corresponding standard deviations. Pearson's correlation coefficient is r = -0.18 with a p-value of 0.65.

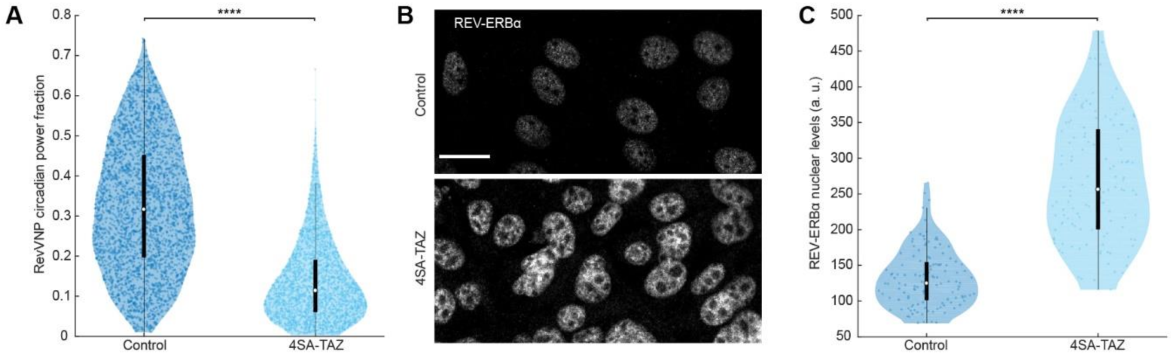


**Figure S5. Overexpression of a constitutively active TAZ mutant affects REV-ERBα expression.**

(A) Violin plots representing the distribution of the single-cell RevVNP circadian power fractions of control and TAZ-overexpressing cells carrying a 4SA dominant positive mutation; n = 3097 and 2359, respectively, from five independent experiments. Two-sided Wilcoxon rank sum test; **** indicates p-value < 0.0001.

(B) Confocal microscopy images of control and 4SA-TAZ cells immunostained against REV-ERBα. All images are displayed under the same brightness and contrast settings. Scale bar, 20 µm.

(C) Violin plots representing the single-cell levels of nuclear REV-ERBα. n = 134 for the control and 123 for 4SA-TAZ, from one representative of three independent experiments. Two-sided Wilcoxon rank sum test; **** indicates p-value < 0.0001.

| Figure 1C | |
|---|---|
| | *High Density* |
| *Low Density* | 6.38E-55 |

| Figure 1E | |
|---|---|
| | *High Density* |
| *Low Density* | 1.59E-287 |

| Figure 2D | |
|---|---|
| | *Conditioned medium* |
| *Control* | 0.4168 |

| Figure 2F | | |
|---|---|---|
| | *Low Density* | *Patterned* |
| *High Density* | 1.83E-21 | 0.031 |
| *Low Density* | N/A | 1.35E-13 |

| Figure 4A | | | |
|---|---|---|---|
| | *5SA-YAP* | *5SA-CCDEL-YAP* | *5SA-S94A-YAP* |
| *Control* | 6.92E-87 | 6.50E-218 | 1.29E-07 |
| *5SA-YAP* | N/A | 6.00E-22 | 3.58E-71 |

| Figure 4D | | | |
|---|---|---|---|
| | *5SA-YAP* | *5SA-CCDEL-YAP* | *5SA-S94A-YAP* |
| *Control* | 1.12E-47 | 2.70E-42 | 2.65E-91 |

| Figure 4E | | | |
|---|---|---|---|
| | *5SA-YAP* | *5SA-CCDEL-YAP* | *5SA-S94A-YAP* |
| *Control* | 9.70E-64 | 4.12E-29 | 2.96E-14 |

**Table S1. P-values of Figures 1 to 4.**
